# Diet induced obesity and type 2 diabetes drives exacerbated sex-associated disease profiles in K18-hACE2-mice challenged with SARS-CoV-2

**DOI:** 10.1101/2022.04.26.489580

**Authors:** Katherine S. Lee, Brynnan P. Russ, Ting Y. Wong, Alexander M. Horspool, Michael T. Winters, Mariette Barbier, Justin R. Bevere, Ivan Martinez, F. Heath Damron, Holly A. Cyphert

## Abstract

SARS-CoV-2 infection results in wide-ranging disease manifestation from asymptomatic to potentially lethal. Infection poses an increased threat of severity to at-risk populations including those with hypertension, diabetes, and obesity. Type 2 Diabetes (T2DM), is characterized, in part, by insulin insensitivity and impaired glucose regulation. T2DM patients have increased disease severity and poorer outcomes with COVID-19. We utilized the diet-induced obesity (DIO) model of Type 2 Diabetes in SARS-CoV-2-susceptible K18-hACE2 transgenic mice to better understand the obesity co-morbidity. Female DIO, but not male DIO mice challenged with SARS-CoV-2 were observed to have shortened time to morbidity compared to normal diet mice. Increase in susceptibility to SARS-CoV2 in female DIO was associated with increased total viral RNA burden compared to male mice. RNAseq analysis was performed on the lungs of non-challenged, challenged, females, males, of either normal diet or DIO cohorts to determine the disease specific transcriptional profiles. DIO female mice had more total activated genes than normal diet mice after challenge; however, male mice experienced a decrease. GO term analysis revealed the DIO condition increased interferon response signatures and interferon gamma production following challenge. Male challenged mice had robust expression of antibody-related genes suggesting antibody producing cell localization in the lung. DIO reduced antibody gene expression in challenged males. Collectively this study establishes a preclinical T2DM/obesity co-morbidity model of COVID-19 in mice where we observed sex and diet specific responses that begin to explain the effects of obesity and diabetes on COVID-19 disease.

## Introduction

Severe Acute Respiratory Syndrome Coronavirus 2 (SARS-CoV-2) continues to pose a worldwide epidemiological threat due to the emergence of novel variants with enhanced transmissibility and disease-causing capabilities. While our understanding of the virus has increased dramatically since its emergence in 2020, questions regarding the mechanisms behind the heterogenous nature of lethality remain. While many infections are asymptomatic or result in mild disease phenotypes, SARS-CoV-2 variants of concern (VOC) remain a considerable threat to at-risk populations where they increase host susceptibility to more severe, deadly infection (1–3). At-risk populations of concern include males, the elderly, pregnant individuals, and those with preexisting conditions including obesity or metabolic disease which make them immunocompromised (2, 4). It is widely accepted that the immune response to pathogen is influenced by biological sex. The distribution of genes related to immunological function across the X and Y chromosomes, contribute to more robust immune response in females that are dampened in males (5, 6). Hormonal differences are thought to suppress the immune system in males while boosting cytokine production in females (7). The compounding effect of physiological factors during the host response to SARS-CoV-2 infection makes identification of exact causative mechanisms difficult. However, increased susceptibility to infections in males and autoimmunity in females have been modeled extensively in preclinical and clinical settings alike (5). It is believed that more deaths from COVID-19 have occurred in men than women making it important to categorize the sex-based differences that occur against SARS-CoV-2 to identify solutions (8).

Another prominent comorbidity, obesity has risen to “epidemic status” in the United States and other countries, occurring at an incidence above 35% in many states (9, 10). Obesity is often associated with the subsequent development of comorbidities akin to those which predispose to severe COVID-19 disease outcomes such as Type 2 Diabetes (11–16). Obesity often occurs concurrent to Metabolic Syndrome, a condition marked by “central” obesity with high adiposity, insulin resistance, and high blood glucose (17–19). Adiposity (increases in adipose tissue distribution) is accompanied by enlargement of individual adipocytes which become stressed and hypoxic at the cellular level. Chronic exposure to stress signals, hypoxic conditions, and oxidative stress causes adipocytes to produce cytokines like CRP, TNF-alpha, and IL-6 in addition to their healthy secretions intended to maintain homeostasis (20). The resulting recruitment to and activation of proinflammatory-type macrophages cells within the adipose tissue raises basal inflammation systemically in a phenomenon known as “metabolic inflammation” (21–23). This inflammation contributes to metabolic dysfunction like insulin resistance in obese persons but also increases susceptibility to pathogens through cellular interactions that are deleterious over time (11–13). COVID-19 also encompasses a complex inflammatory milieu where delayed interferon responses allow virus to continue replicating, meanwhile inciting the proinflammatory actions of neutrophils and lymphocytes that drive the disease-characteristic “cytokine storm” (17). The contributions of preexisting metabolic dysfunction to this aberrant inflammatory response have yet to be mechanistically defined. In laboratory mice, severe outcomes for obese individuals during infection have been modeled for numerous agents including influenza, West Nile Virus, and even the parasite *Leishmania* (24–26). Similar studies have implicated preclinical diabetes models as comorbid conditions, but no extensive work has been done to define the comorbid outcomes of SARS-CoV-2 (27).

T2DM is estimated to be the second-most common comorbidity in patients with severe COVID-19, resulting in a 2-3 times greater likelihood to succumb compared to healthy persons (28). Over 460 million people worldwide have been diagnosed with diabetes mellitus (either T1DM or T2DM) and greater than 60% of type 2 diabetics are also clinically characterized as obese (29). SARS-CoV-2 infection combined with the metabolic dysfunction in T2DM is associated with an increased risk of pneumonia requiring ventilation, ICU admission, and “long COVID” (30). While COVID-19 vaccine implementation around the world has been a positive effort for protecting vulnerable populations, T2DM has been linked to reduced COVID-19 vaccine efficacy, with lower IgG and neutralizing antibody development (14, 31). Because of their predisposed risk to severe outcomes, defining the immunological profile of type 2 diabetics is a necessary step towards solving vaccine-established protection discrepancies.

Most of our knowledge regarding the positive correlation between metabolic dysfunction and SARS-CoV-2 severity comes from retrospective clinical studies, where it becomes impossible to discern the molecular mechanisms that governed severe outcomes (32–40). In order to identify and characterize the mechanisms behind increased infection and severity we developed a preclinical model of disease comorbidities using the K18-hACE2 transgenic mouse model and diet induced obesity (DIO) where metabolic disease is confirmed by the development of T2DM (41,42,51,52,43–50). K18-hACE2 mice were subjected to the high-fat, high carbohydrate diet for 8 weeks causing measurable obesity, metabolic dysfunction, and hyperglycemia. Normal diet and DIO males and females were either mock challenged or challenged with the Alpha variant of SARS-CoV-2. Female DIO mice were observed to have shorter time to morbidity than normal diet mice while DIO diet female mice also exhibited higher viral RNA burden at the time of terminal euthanasia, indicating sex differences in disease pathology. RNAseq analysis was used to characterize the transcriptional responses of the lung in all experimental cohorts. Systems based analysis revealed DIO mice have unique responses to SARS-CoV-2 challenge including lack of antibody-related gene diversity compared to normal diet K18-hACE2 mice in addition to differential gene expression profiles. Our data illustrate how metabolic dysfunction can enhance COVID-19 disease and suggest a synergism between hyperglycemia and gene expression profile changes. This data helps to link molecular alterations with infection severity, thus constructing a profile of potential therapeutic targets for the treatment and prevention of death by COVID-19 illness.

## Materials and methods

### Animal, Ethics, Biosafety statement

All research performed was approved by West Virginia University IACUC protocol number 2004034204. K18-hACE2-mice (B6.Cg-Tg(K18-ACE2)2Prlmn/J; JAX strain number #034860). All SARS-CoV-2 viral propagation or challenge studies were conducted in the West Virginia University Biosafety Laboratory Level 3 facility under the IBC protocol number 20-04-01. SARS-CoV-2 infected mouse serum and lung supernatants were inactivated with 1% Triton per volume before exiting high containment. Additional tissues were treated using TRIzol reagent (Zymo Research Catalog No R2050-1-200) at a ratio of at least 1:1 or fixed with 10% neutral-buffered formalin before additional work in BSL2 conditions.

### High fat diet induced obesity and Type 2 Diabetes model and intraperitoneal glucose tolerance test (IPGTT)

Diet induction of obesity was achieved through feeding a high fat diet (Bio-serv Mouse diet high fat 60% kCAL from fat #S3282) for 8 weeks to cohorts of 6-week-old female and male K18-hACE2-mice. Concurrently, control age-matched K18-hACE2-mice remained on a standard chow diet. Weight was monitored weekly. Intraperitoneal glucose tolerance testing (IPGTT) was performed after the 8-week diet induction period on mice fasted for 6h. Tail blood was collected prior to intraperitoneal injection of glucose as the baseline (Time 0 minutes). Glucose was injected at 2mg/g body weight prepared in sterile PBS (20% w/v). Blood glucose was measured at 30 minutes and 60 minutes in blood collected from a tail snip using a hand-held glucometer

### SARS-CoV-2 cultivation and K18-hACE2 mouse challenge

The SARS-CoV-2 Alpha variant strain was obtained for challenge from BEI: hCoV19/England/204820464/2020 (Alpha; NR-54000)(GISAID: EPI_ISL_683466). The virus was propagated in Vero E6 cells (ATCC-CRL-1586) as described previously (53). Normal diet or DIO K18-hACE2 mice were challenged with a 10^3^ PFU dose. Viral doses were prepared from the first passage collections from infected Vero E6 cells. Mice were anesthetized with an intraperitoneal injection of ketamine (Patterson Veterinary 07-803-6637) / xylazine (Patterson Veterinary 07-808-1947) at a concentration of 80 mg/kg. K18-hACE2 mice were challenged with virus by intranasal administration of 25μL dose per nare (50μL total).

### Disease scoring of SARS-CoV-2 challenged mice

Challenged K18-hACE2 mice were evaluated daily through in-person health assessments in the BSL3 facility as well as surveillance using SwifTAG Systems video monitoring. Health assessments of the mice were scored based on the following criteria: weight loss (scale 0-5 (up to 20% weight loss)), appearance (scale 0-2), activity (scale 0-3), eye closure (scale 0-2), and respiration (scale 0-2). All five criteria were scored based off a scaling system where 0 represents no symptoms and the highest number on the scale denotes the most severe phenotype as previously described by our lab (54). Additive health scores of the criteria listed above were assigned to each mouse after evaluation and assessed so that mice scoring 5 or above, 20% weight loss, or significant drops in temperature received immediate euthanasia. Cumulative disease scoring was calculated by adding the disease scores of each mouse within the group on each day. Morbid mice that were euthanized during the study before day 14, retained their disease score for the remainder of the experiment for reporting purposes.

### Euthanasia and tissue collection

Mice were euthanized either due to disease scores or at the end of the experiment with an IP injection of Euthasol (390mg/kg) (Pentobarbital) followed by cardiac puncture as a secondary measure of euthanasia. Each animal was the dissected to collect tissues for pathological analysis as previously described (55)Cardiac blood was collected in BD Microtainer gold serum separator tubes and centrifuged at 15,000 x *g* for 5 minutes to separate the serum for downstream analysis. PBS (1mL) was pushed by catheter through the nasal pharynx and collected in a 1.5mL Eppendorf tube for Nasal Wash. For future RNA purification, 500μL of nasal wash was added to 500μL of TRIzol reagent (Zymo Research Catalog No R2050-1-200) and the remainder of the nasal wash was stored for serological analysis. Lung and brain tissues were dissected from each animal. The right lobe of the lung was homogenized in 1mL PBS in gentleMACS C tubes (order number: 130-096-334) using the m_lung_02 program on the gentleMACS Dissociator. For RNA purification from the tissues, lung homogenate (300μL) was added to 1000μL of TRIzol Reagent. For serology and cytokine analysis, 300μL of lung homogenate was centrifuged at 15,000 x *g* for 5 minutes to separate and collect the supernatant. Brain tissue was also homogenized in 1mL PBS using gentleMACS C tubes and the same setting as lung on the gentleMACS Dissociator. From the homogenate, 500 μL was combined with 1000μL of TRI Reagent for RNA purification.

### qPCR SARS-CoV-2 viral copy number analysis of lung, brain, and nasal wash

From the lung, brain and nasal wash, RNA was purified using the Direct-zol RNA miniprep kit (Zymo Research R2053) and the manufacturer’s protocol. qPCR using the Applied Biosystems TaqMan RNA to CT One Step Kit (Ref: 4392938) was performed to quantify SARS-CoV-2 copies. Winkler. *et al*, 2020 (56) methods were used to synthesize the nucleocapsid primers (F: ATGCTGCAATCGTGCTACAA; R: GACTGCCGCCTCTGCTC) and TaqMan probe (IDT:/56-FAM/TCAAGGAAC/ZEN/AACATTGCCAA/3IABkFQ/). According to the Applied Biosystems TaqMan RNA to CT One Step Kit manufacturer protocol, 2XTaqMan RT-PCR Mix, 900nM Forward and reverse primers, 250nM TaqMan probe, 40X TaqMan RT enzyme mix and 100ng RNA template, were combined for each reaction. If sample RNA was purified at a concentration less than 100ng/uL they were not diluted for the qPCR reaction. Each sample was prepared in triplicate and loaded into a MicroAmp Fast optical 96 well reaction plate (Applied Biosystems 4306737). The plates were run on the StepOnePlus Real-Time System machine using the parameters: reverse transcription for 15 minutes at 48°C, activation of AmpliTaq Gold DNA polymerase for 10 minutes at 95°C, 50 cycles of denaturing for 15 seconds at 95°C, annealing at 60°C for 1 minute.

### Serological analysis

ELISA assays were performed as previously described (57) coating high binding plates overnight with either 2μg/mL nucleocapsid or 2μg/mL RBD. After sample was added, either goat-anti-mouse secondary IgG HRP (1:2000 dilution) to measure IgG or goat-anti-mouse IgM HRP (1:10000 dilution) to measure IgM was used.

### Cytokine analysis

To measure IFN-γ as well as other cytokines, samples of lung supernatant from each mouse were run on the R&D 5-plex mouse magnetic Luminex assay (Ref LXSAMSM). The manufacturer’s protocols were followed to prepare and run samples. The plate was analyzed on the Luminex Magpix to calculate concentrations (pg/mL) based off of the individual standard curves for each cytokine. MSD assay plates were analyzed using the Meso Scale Discovery Sector 2400.

### Illumina library preparation, sequencing, and *in silico* bioinformatic analysis

RNA concentrations were measured with the Qubit 3.0 Fluormeter using the RNA high sensitivity kit (Life Technologies) and RNA integrity was assessed using an Agilent TapeStation. RNA was DNAased before library preparation. Illumina sequencing libraries were created with the KAPA RNA HyperPrep Kit with RiboErase (Basel, Switzerland). Resulting libraries passed standard Illumina quality control PCR and were sequenced on an Illumina NovaSeq s4 4000 at Admera Health (South Plainfield, NJ). A total of ∼100 million 150 base pair reads were acquired per sample. Sequencing data will be deposited to the Sequence Read Archive. The reads were trimmed for quality and mapped to the *Mus musculus* reference genome using CLC Genomics Version 21.0.5. An exported gene expression browser table is provided as supplemental materials (Table S1). Statistical analysis was performed with the Differential Expression for RNA Seq tool and genes were annotated with the reference mouse gene ontology terms. PCA plots were formed in CLC Genomics Version 21.0.5. Quantification of the number of activated or repressed genes unique to each experimental group was performed using Venny 2.1 (58) and visually modelled using the WEB-based BioVenn (59). Genes from each experimental comparison with significant fold changes compared to no-challenge (Bonferroni ≤ 0.04) were submitted to the WEB-based Gene SeT AnaLysis Toolkit’s Over Representation Analysis (ORA) software compared to the reference set “affy mg u74a” to determine GO terms from gene ontology and biological process databases (FDR ≤ 0.05) (60). GO Term heat maps were generated using Morpheus (61). In order to analyze the expression of the hACE2 transgene, the RNA reads were mapped to the human ACE2 gene (GRCh38). SARS-CoV-2 reads were analyzed by mapping the reads to the SARS-CoV-2 WA-1 reference genome. hACE2 and viral reads were normalized by dividing counts per 50M total reads in each sample. Raw read data is available at NCBI SRA: SUBXXXXXX (submission complete, pending processing).

### Ingenuity Pathway Analysis

RNAseq fold change gene expression data was submitted to Ingenuity Pathway analysis using a cut off of *P=* 0.05. Pathways that were statistically enriched were exported and plotted into heat maps using Morpheus as described above.

### Statistical analyses

Tests to determine statistical significance were performed using GraphPad Prism version 9. In all DIO K18-hACE2 mouse studies n = 5 per group. In challenge dose determination studies, n ≥ 3. Statistically significant differences between Kaplan-Meyer curves were analyzed using Mantel-Cox log-rank tests. Student’s t-tests were used for comparisons made between two groups. When three or four groups were being compared, statistical differences were assessed using one-way ANOVA with Dunnett’s multiple comparisons test or Two-Way ANOVA with Tukey’s multiple comparisons test for parametric data. For any non-parametric data, Kruskal-Wallis tests with Dunn’s multiple comparisons tests were used.

## Results

### K18-hACE2 mice develop obesity, metabolic dysfunction, and type two diabetes due to high fat diet

The COVID-19 pandemic has illustrated that humans respond to infection with a great deal of heterogeneity. SARS-CoV-2 infection can be lethal in some patients but cause mild or asymptomatic disease in others. Due to this heterogeneity in infection severity, it is important to understand co-morbidities in order to develop therapeutic interventions that support the most at-risk populations. To understand the impact of metabolic dysfunction as seen in obesity and Type 2 Diabetes on the outcomes of viral infection, we utilized a diet-induction model of obesity with K18-hACE2 transgenic mice (Fig. 1A). Compared to normal chow mice, DIO mice gained 25% or 37% bodyweight, in females and males respectively (Fig. 1B). Intraperitoneal glucose tolerance testing after 8-weeks of a high-fat or normal diet, was utilized to evaluate their ability to clear glucose. Glucose tolerance was impaired significantly in male mice fed the DIO diet (*P<*0.0001) while female mice presented with non-significant low to mild impairment (Figure 1C-D). The DIO model has been previously used to study the effects of obesity and Type 2 Diabetes in mice, and our data here suggest that K18-hACE2-mice on the DIO diet do develop metabolic dysfunction (51, 52).

**Figure 1:**
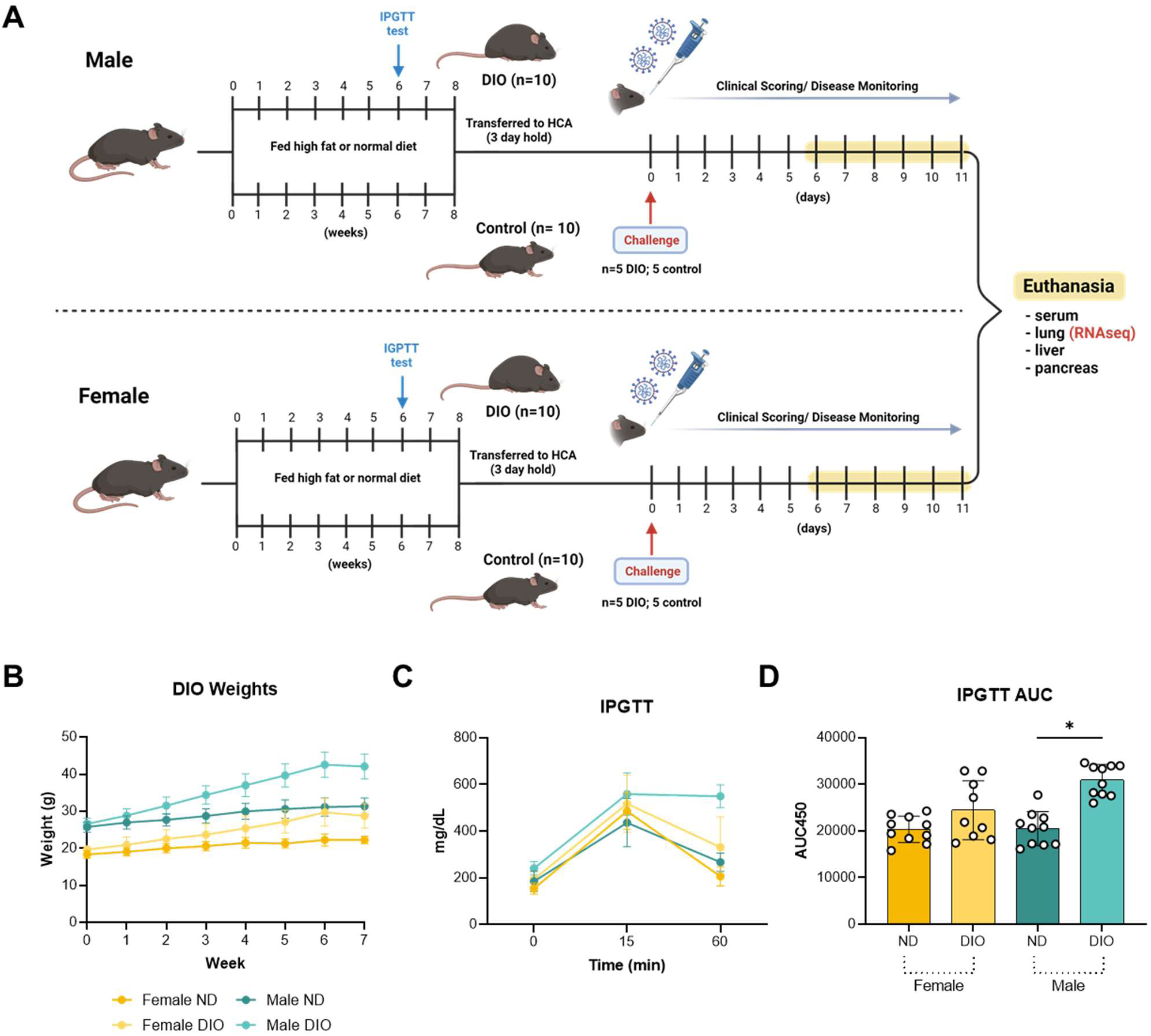
Development of the Diet Induced Obesity (DIO) K18-hACE2 transgenic mouse model. Male and female K18-hACE2 transgenic mice were given a high-fat, high-carbohydrate diet for 8 weeks to induce glucose impairment consistent with the Diet Induced Obesity model (n=20 males; 20 females) before challenge with 10^3^ PFU SARS-CoV-2 Alpha variant. Control groups that were age- and sex-matched remained on normal diet for the entirety of the experiment (n=10 DIO and 10 normal diet males; 10 DIO and 10 normal diet females) (A). DIO as well as normal-diet mice were weighed weekly to measure DIO progression (*P=* 0.0207 female ND vs Female DIO; *P=*0.0281 Male ND vs ale DIO) (B). At week six of the high-fat, high-carbohydrate diet, DIO and normal-diet mice underwent intraperitoneal glucose tolerance testing (IGPTT) which revealed impaired glucose clearance and fasting hyperglycemia in male DIO mice (C,D) (*P<0.0001* Male ND vs Male DIO; *P=*NS Female ND vs Female DIO).

### DIO shortens the time to morbidity in lethal SARS-CoV-2 challenge

The Alpha variant (strain B.1.1.7) of SARS-CoV-2 emerged in the United Kingdom in the spring of 2021 and moved rapidly across the globe. Alpha seemed to have enhanced virulence compared to ancestral strains of the virus in K18-hACE2-mice as well as other preclinical animal models (55,62–67). Due to the significance of Alpha variant’s dominance among circulating strains in humans, we used this VOC to study the effects of DIO and Type 2 Diabetes in the K18-hACE2 mouse model of SARS-CoV-2 infection. Prior to the experiment, optimization of viral challenge dose was performed to determine the dose for achieving symptomatic disease (Fig. 2A-B). The Alpha strain was intranasally administered at 10^3^, 10^4^, and 10^5^ PFU per dose. Using a previously established disease scoring system (55,68,69), we determined that the 10^3^ PFU dose caused lethality but postponed the time to morbidity compared to higher doses (Fig. 2A) and caused disease phenotypes that increase in severity over time (Fig. 2B). Male and female mice were utilized for DIO challenge studies to account for sex-based predispositions to disease severity (70, 71). DIO SARS-CoV-2 challenged mice experienced a shortened time to morbidity in both females and males compared to their normal-diet controls. Female DIO mice experienced 0% survival at 7 days post challenge (median time to death=6 days) compared to only 20% survival in female controls (median time to death=8 days) (Fig. 2C). DIO and normal-diet male mice responded to challenge in a similar manner, with no statistical differences in survival (median time to death=6 days in both groups). Daily disease scoring over the post-challenge period trended in a similar pattern in DIO and control challenged mice with no sex-dependent differences (Fig. 2E-F). Collectively, these data suggest that the DIO condition has a greater impact on survival in female mice suggesting that sex specific responses may impact COVID-19 pathogenesis.

**Figure 2:**
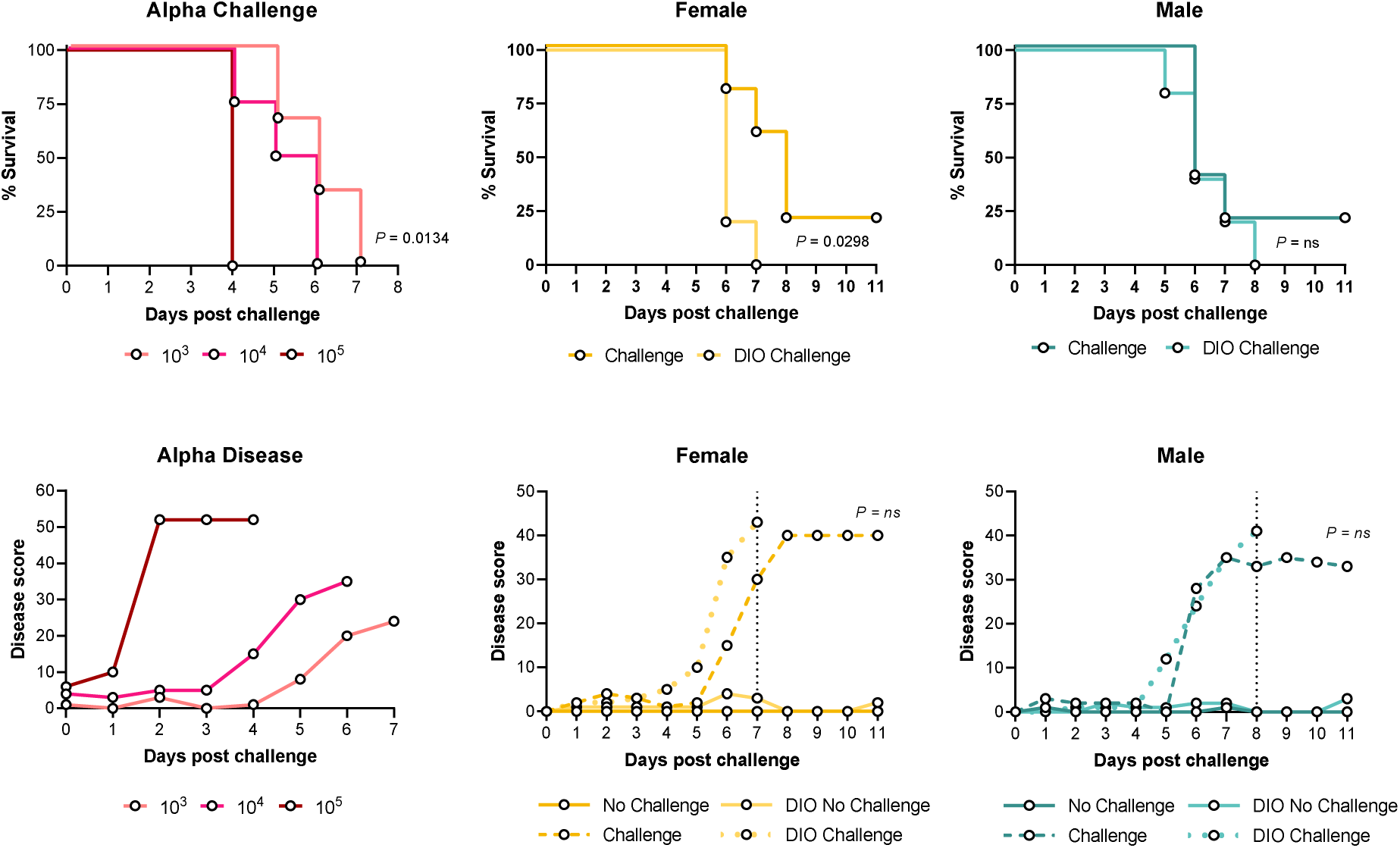
Survival and disease scores of SARS-CoV-2 challenged DIO-K18-hACE2 mice. An intranasal challenge dose of 10^3^ PFU Alpha SARS-CoV-2 is sufficient for causing disease phenotypes with delayed morbidity in K18-hACE2 mice (10^3^ PFU n=3; 10^4^ PFU n=4; 10^5^ PFU n=5)(A,B). Survival and disease scores post-challenge were measured for female DIO and normal-diet mice (C,D), as well as male DIO and normal-diet K18-hACE2 mice (E,F). Log-Rank (Mantel-Cox) tests were utilized to measure the significance level of changes in survival curves. Unpaired T-tests were used to test the significance of disease scores between normal diet and DIO challenge groups. Dotted lines indicate day at which full morbidity of group had been achieved. n=10 in panels C, D, E, F. ns= no significance

### Obese female mice experience greater viral RNA burden in the lungs

To begin identifying the factors that may contribute to the changes in survival that were observed in DIO mice, we next investigated differences in viral burden measured by qRT-PCR analysis of lung tissue for nucleocapsid transcript copy number. Total lung RNA was isolated from mice at euthanasia at their respective humane endpoints (Fig. 2CD). Viral RNA burden was found to be higher in the lungs of DIO female mice than in female normal diet controls and no difference was seen in the viral RNA burden of males (Fig. 3A). Normal diet female mice were observed to have approximately 1 million copies of nucleocapsid RNA transcripts on average per lung lobe whereas DIO females have 100-fold more copies, suggesting DIO enhances viral burden in females (Fig. 3A). The enhanced viral burden, per PCR analysis, suggested that the differences in time to morbidity and survival in females (Fig. 2C), may be related to viral burden or inability to clear virus.

**Figure 3:**
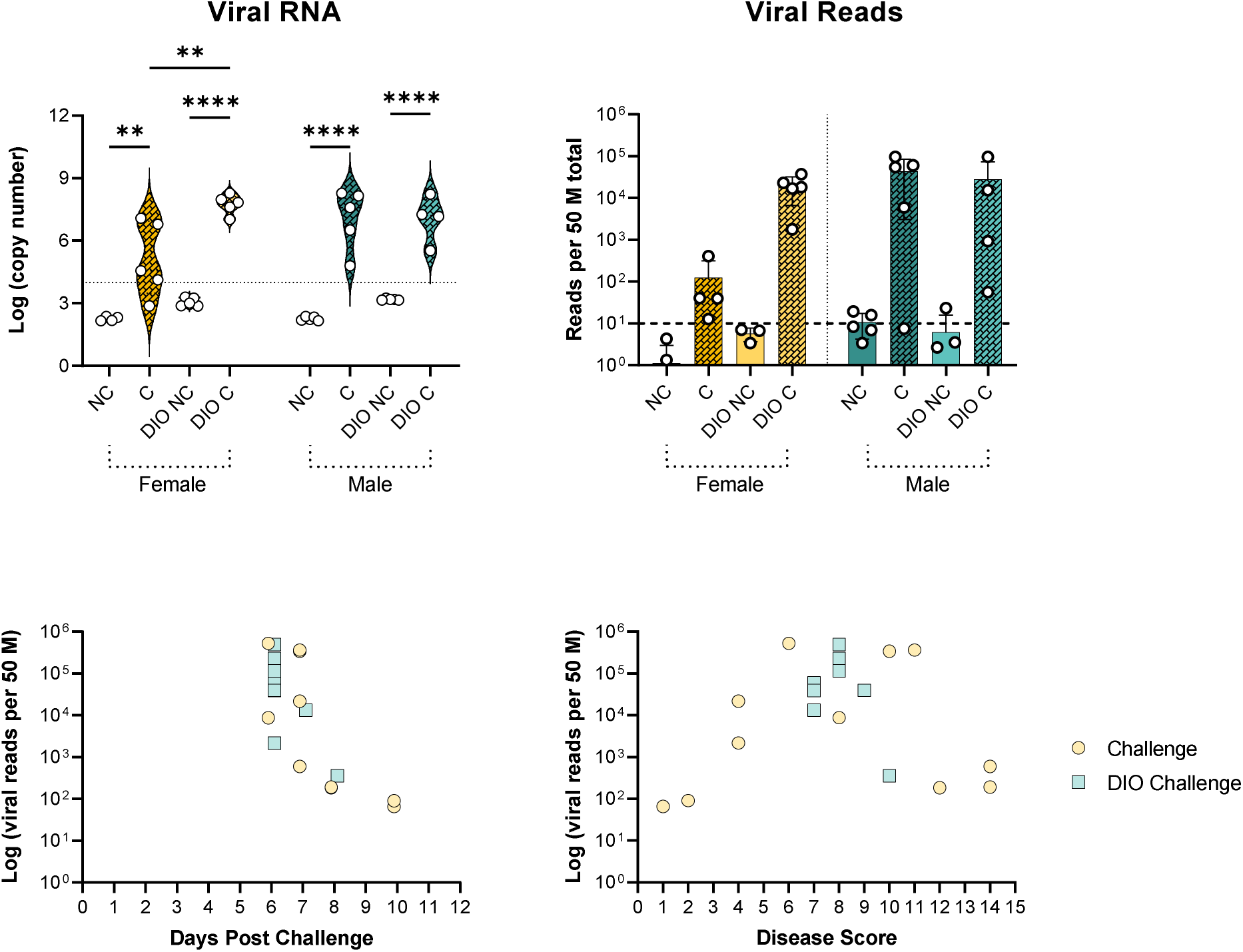
Quantification of viral burden in the lung by qPCR and RNAseq. Viral nucleocapsid RNA was detectable via qPCR in the lung tissue of challenged- and DIO-challenged K18hACE2 mice (one-way ANOVA ** P=0.0020; ****P=<0.0001)(A). SARS-CoV-2 nucleocapsid reads were additionally quantified using RNAseq of lung RNA samples (ns)(B). In order to identify correlations between viral RNA burden and disease severity, RNAseq nucleocapsid reads were plotted against host’s euthanasia day (C) and individual disease score at euthanasia (D). Dotted line indicates limit of detection via qPCR. Dashed line indicates number of RNAseq reads that were examined and determined to be nonspecific. NC=Normal Diet No Challenge; C=Normal Diet Challenge; DIO NC=DIO No Challenge; DIO C=DIO Challenge.

### Transcriptomic analysis of viral RNA confirms females have increased viral RNA due to DIO condition

As a secondary method of evaluating viral burden at, total lung RNA was used to perform bulk RNAseq analysis to measure the number of virus gene transcripts per total tissue RNA. Viral RNA reads were mapped to the SARS-CoV-2 reference genome and represented per 50M illumina reads obtained per sample. Total viral reads perfectly mirrored the nucleocapsid qRT-PCR analysis (Fig. 3B). A 100-fold increase in viral RNA was also observed for DIO females compared to normal females (Fig. 3B). To identify correlations between viral burden and morbidity, total viral reads were plotted against the day post-challenge that humane euthanasia occurred (Fig. 3C) or against disease score (Fig. 3D). High viral reads corresponding to the mortality window 6 days post-challenge were observed for DIO mice where normal diet mice which survived longer underwent euthanasia (due to disease score or planned experimental endpoint) at lower viral burdens. Disease scoring comparisons had greater variance, but DIO mice trended towards having higher viral reads in the lung with higher disease scores (Fig. 3D).

### Transcriptomic analysis of mouse gene expression profiles unique to metabolic dysfunction

Bulk RNAseq analysis of infected mouse tissues allows for simultaneous pathogen and host transcriptomic analysis (72). We have previously used similar techniques to characterize mouse and bacterial gene expression during infection (73–75). After evaluation of the viral transcriptomics of SARS-CoV-2 challenge in DIO mice, lung tissue RNA from non-challenged, challenged, DIO, or normal diet mice was used to characterize the host transcriptomic responses to SARS-CoV-2. Basic gene expression profiles of biological replicates in the experimental groups were compared using Principal Component Analysis. The transcriptional profiles showed distinct patterns of gene expression between challenge and no challenge groups. Interestingly, we observed a higher overlap between DIO and normal-diet in males and female mice than between challenged and non-challenged mice regardless of diet (Fig. 4A and B). This suggested that challenge with SARS-CoV-2 has a greater impact on the lung transcriptome than diet in K18-hACE2 mice. Separation of the gene profiles into activated and repressed expression bins allowed for visualization of sex-driven differences. DIO induction increased the number of uniquely activated and repressed genes upon viral challenge in females but decreased it for males (Fig. 4C). Male DIO mice had smaller unique transcriptional profiles (589 DIO male-specific genes) while female DIO mice had a much larger transcriptional response (1974 DIO female-specific genes). When the pools of activated genes (*P* < 0.05) from each experimental group were compared using a Venn Diagram, the unique gene profiles became narrower, and the unique expression profiles stemming from sex or DIO induction could be appreciated (Fig. 4D). While a core set of 835 genes were found to be activated in all SARS-CoV-2 challenged mice, DIO led to unique transcriptional profiles with differential expression of 765 unique genes in DIO females and 415 unique genes in DIO males. These data suggest that DIO and sex both influence the response to SARS-CoV-2 challenge.

**Figure 4:**
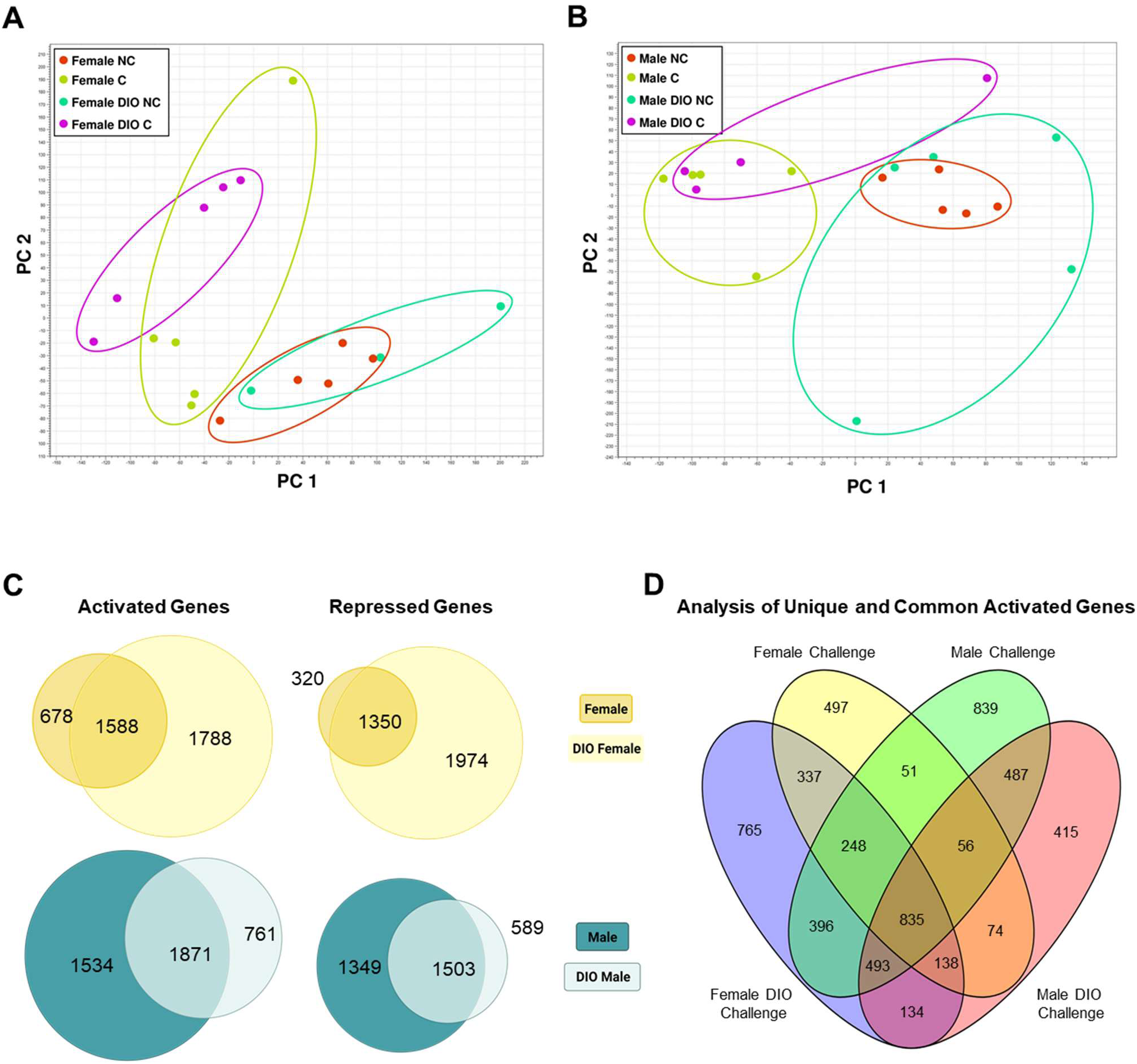
Transcriptomic analysis of lung tissues from Alpha SARS-CoV-2 challenged DIO and normal diet mice. Principal component analysis was used to compare differential gene expression profiles of viral challenged DIO or control female (A) and male mice (B). RNAseq reads from challenged DIO or normal diet mice were compared to no-challenge lean mice to determine relative gene activation or repression (C). Venn Diagrams were used to visualize the unique expression of genes that were specific to experimental condition.

### Ingenuity pathway analysis identifies unique canonical pathway matches from metabolic dysfunction and infection

RNAseq analysis showed that the added variables of sex and preexisting comorbidities change the host transcriptional response to SARS-CoV-2 challenge by changing the unique differential gene expression profiles of K18-hACE2 mice. Gene expression fold change data comparing the profiles of each experimental group to no-challenge normal-diet control mice were evaluated by Ingenuity Pathway Analysis to identify affected canonical pathways suggested by gene expression. The pathways that were associated with greater positive z-scores in the female DIO challenge datasets implicated activated inflammatory signaling pathways and immune cell activation that were not supported by the expression profiles of DIO no-challenge mice (Fig. 5). Male mice’s gene expression profiles had altogether similar canonical pathway associations to females. The addition of the DIO condition to challenge groups augmented but did not significantly vary the z-scores of many pathways. Of note, we observed a decreased association of genes within the T cell receptor signaling pathway in male DIO mice. T cell responses are a major contributor to the antiviral response and are heavily implicated in the host response to COVID-19. The magnitude and polyfunctionality of the T cell response in severe cases of COVID-19 is a predictor of outcome as well as the memory response that is protective against reinfection (76, 77). Dysregulation of the T cell response early on due to comorbidities may be partially responsible for disease outcome.

**Figure 5:**
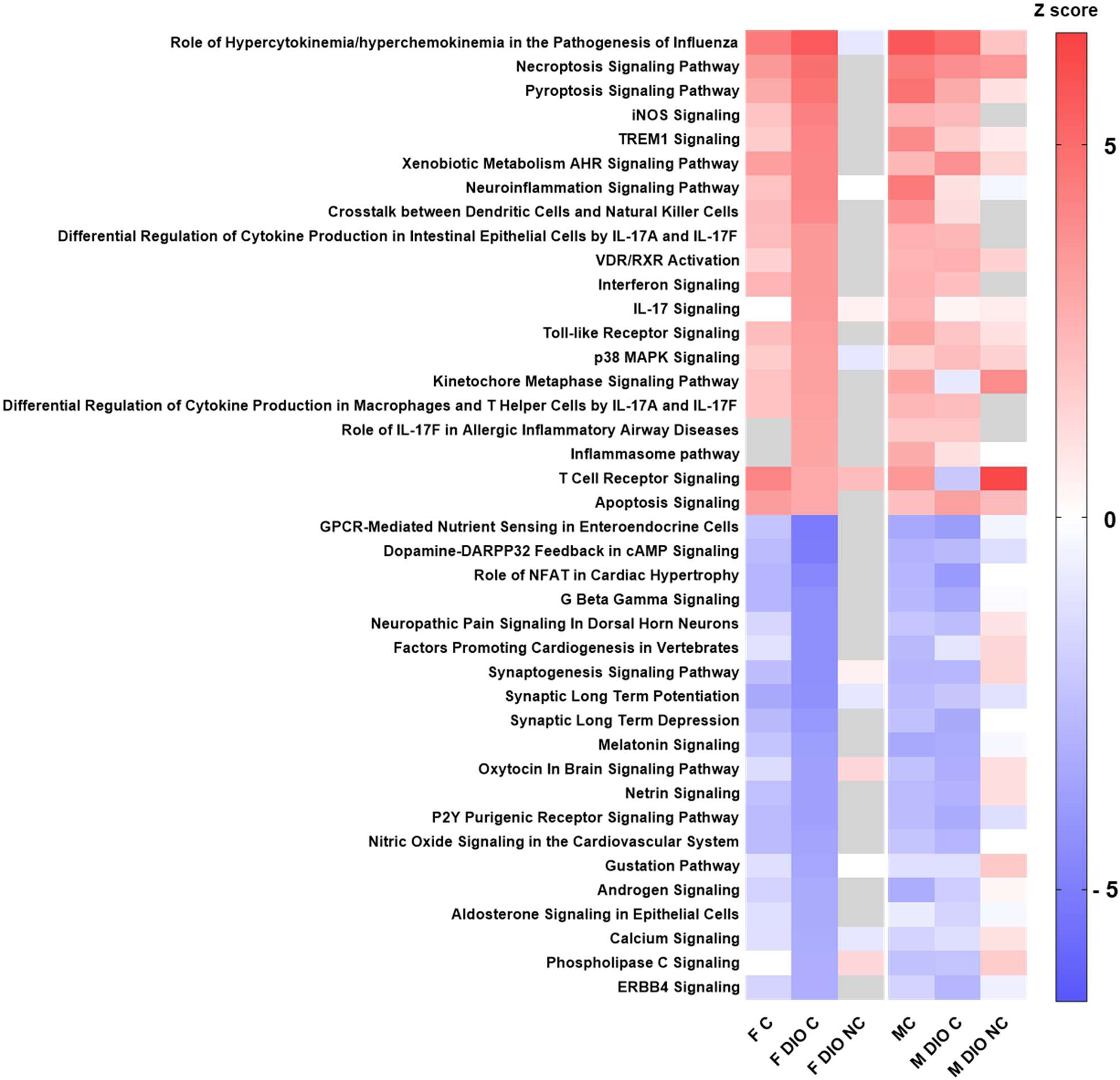
Ingenuity Pathway Analysis of differential gene expression profiles. Fold change values of genes in experimental groups compared to no-challenge were inputted to Ingenuity Pathway Analysis software and sorted into canonical pathways. The highest and lowest 20 canonical pathways by z-score in the female DIO challenge group are shown.

We continue our investigation of differential gene expression profiles by narrowing in on specific IPA pathways. To gain preliminary insights into the changes in T cell responses within the different treatment groups, a heatmap of fold changes (compared to sex-matched normal-diet no-challenge groups) in the expression of genes within the T Cell Receptor Signaling pathway was generated. Interestingly, male DIO mice had higher expression of numerous T cell-related signaling genes, including many that code for the T cell receptor alpha variable region, without SARS-COV-2 challenge that were low in the male DIO challenge group and both groups of female mice (Fig. 6A). Genes like RelA, which encodes the p65 transcription factor for NF-κB signaling (78), and MAPK13, which encodes the pro-inflammatory p38 MAP kinase (79) were upregulated in both male and female DIO challenge groups compared to their normal-diet counterparts. However, genes like PDK1 (80) and CARD11 (81) which support T cell proliferation were high in both normal-diet challenge mice, and comparatively low in both DIO challenge groups. These data suggest DIO resulted in changes in T cell activation. Another pathway of interest was the Coronavirus pathway, which is comprised of known biomarkers that are either activated or repressed during SARS-CoV-2 infection (82). Contrary to the most differential T cell related gene expression occurring in no-challenge DIO males, a large number of inflammatory genes appeared to be up-regulated in challenged DIO females yet repressed in normal-diet females (Fig. 6B). NLRP3, encoding for the antiviral inflammasome, was up in DIO females after challenge compared to all other groups, as were genes for the proinflammatory mediators STAT3 and CCL2 (83, 84). It is possible that the gene expression differences that are associated with these pathways are directly involved in the impaired viral clearance and affected disease pathogenesis experienced due to the DIO condition.

**Figure 6:**
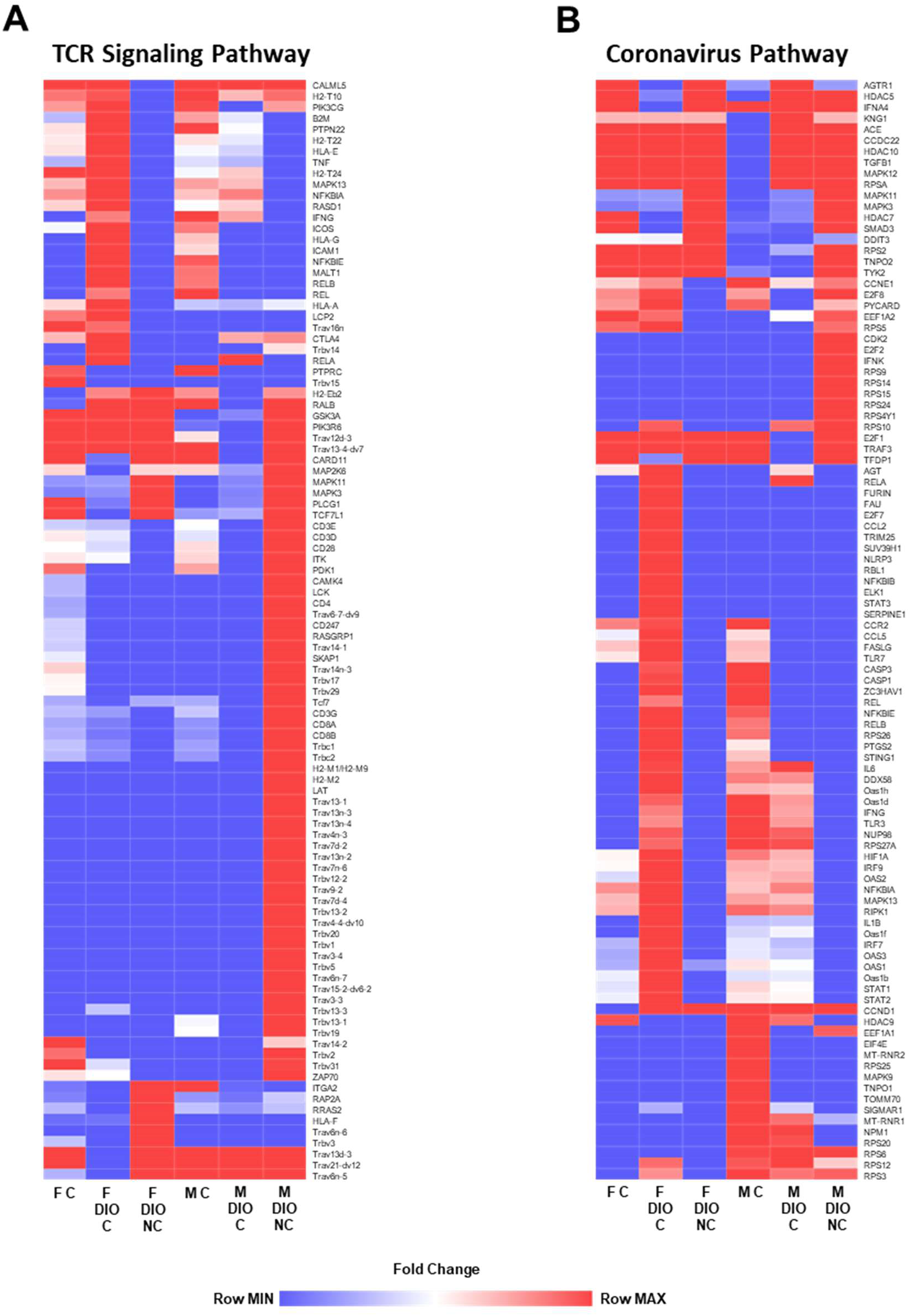
Differential expression of canonical pathways. Heat maps showing the fold change compared to no-challenge of 100 genes in lung RNA from DIO or normal diet mice challenged with SARS-CoV-2 or not within the IPA pathways “T Cell Receptor Signaling” (A) and “Coronavirus Pathogenesis Pathway” (B).

### Metabolic dysfunction in female K18-hACE2 mice heightens host antiviral response profile

To further refine our RNAseq data analysis, Geno Ontology “GO Term” analysis was performed to identify biological processes that were affected in our experiment based on the lung tissues’ transcriptional profiles. Although DIO induction and sex resulted in different lists of suggested GO terms, a conservative list of terms was present in each analysis that was related to activation of the immune response in viral infection. The terms were graphed with their corresponding enrichment ratio to compare their relevance in the genetic profiles of each experimental condition (Fig. 7A). The highest enrichment ratios were seen for the GO terms related to interferon response and were increased in the female DIO challenge sets compared to others. GO terms related to antigen processing and presentation pathways appeared to be absent in both DIO groups. To measure the inflammatory response to virus, of the levels of innate cytokines were measured in serum collected at euthanasia (data not shown). The most striking differences in cytokine production were noted for IFN-γ (Fig. 7B). In females, DIO induction caused an increase in production of IFN-γ compared to minimal production in female challenge control mice (Fig. 7B). DIO induction in males did not result in a significant change in IFN-γ production. This is important as IFN-γ production is triggered during SARS-CoV2 infection and is essential for viral clearance and resolution of the infection(85). Together, these data suggest that changes in the immune response triggered by T2DM/obesity may hinder positive disease outcomes. As shown above, challenged DIO females had 100-fold higher viral RNA burden in the lung (Fig. 3A) and we speculate that the presence of higher levels of viral particles in the lung of DIO females is associated with the increased interferon response observed here.

**Figure 7:**
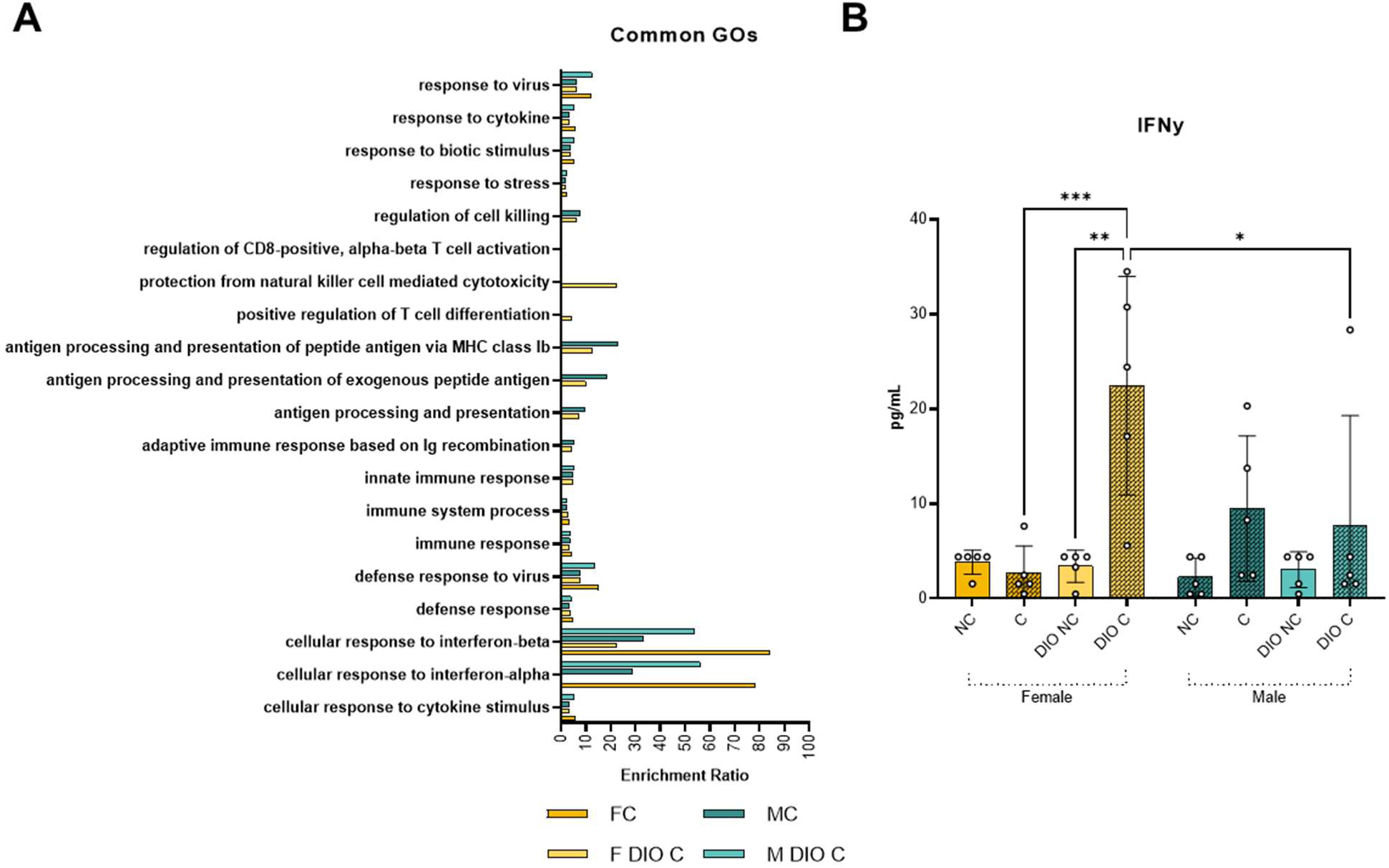
Antiviral response profile of DIO SARS-CoV-2 challenged K18-hACE2 mice. GO terms related to the immune response are graphed to compare term enrichment in experimental groups (A). Interferon gamma was measured in the lung supernatant of K18-hACE2 mice at euthanasia (B). (*P* value: * = 0.0234; ** = 0.0015; *** = 0.0010)

### Sex and metabolic dysfunction influence antibody production

In addition to T cell and interferon response, antibodies also play an important role in the immune response against SARS-CoV-2. RNAseq reads from lung RNA were analyzed to quantify antibody-related gene expression and gain insights into the effects of diet and sex on the humoral response. Overall, a trend was appreciable where antibody genes were highest expressed by male mice challenged with SARS-CoV-2 when reads were mapped visualized by heatmap (Fig. 8A). In male challenged mice, the diversity of upregulated Igh and Igk genes was greater than in the other groups, suggesting: 1) the presence of B cells in the lung, and 2) the unique activation of these B cells in male normal-diet mice after challenge compared to DIO males and both female groups. To determine if changes observed at the mRNA expression level translated into variation in protein antibody responses, quantification of anti-SARS-CoV-2 RBD and anti-nucleocapsid IgG and IgM in serum was performed via ELISA to measure virus-specific antibodies. At this early time point after challenge, only DIO challenged females produced significant levels of anti-RBD IgM antibodies compared to non-challenged female mice. We did not observe statistically significant increases in anti-nucleocapsid IgM, anti-RBD IgG or anti-nucleocapsid IgG in any of the groups, likely due to the fact that titers were measured 11 or fewer days post-challenge. Altogether, lung mRNA and circulating serological immunoglobulin data suggest that the humoral response generated by males and females in response to SARS-CoV2 is different, with the female response characterized by higher levels of circulating IgM and the male response characterized by B cell activation and differentiation in the lung.

**Figure 8:**
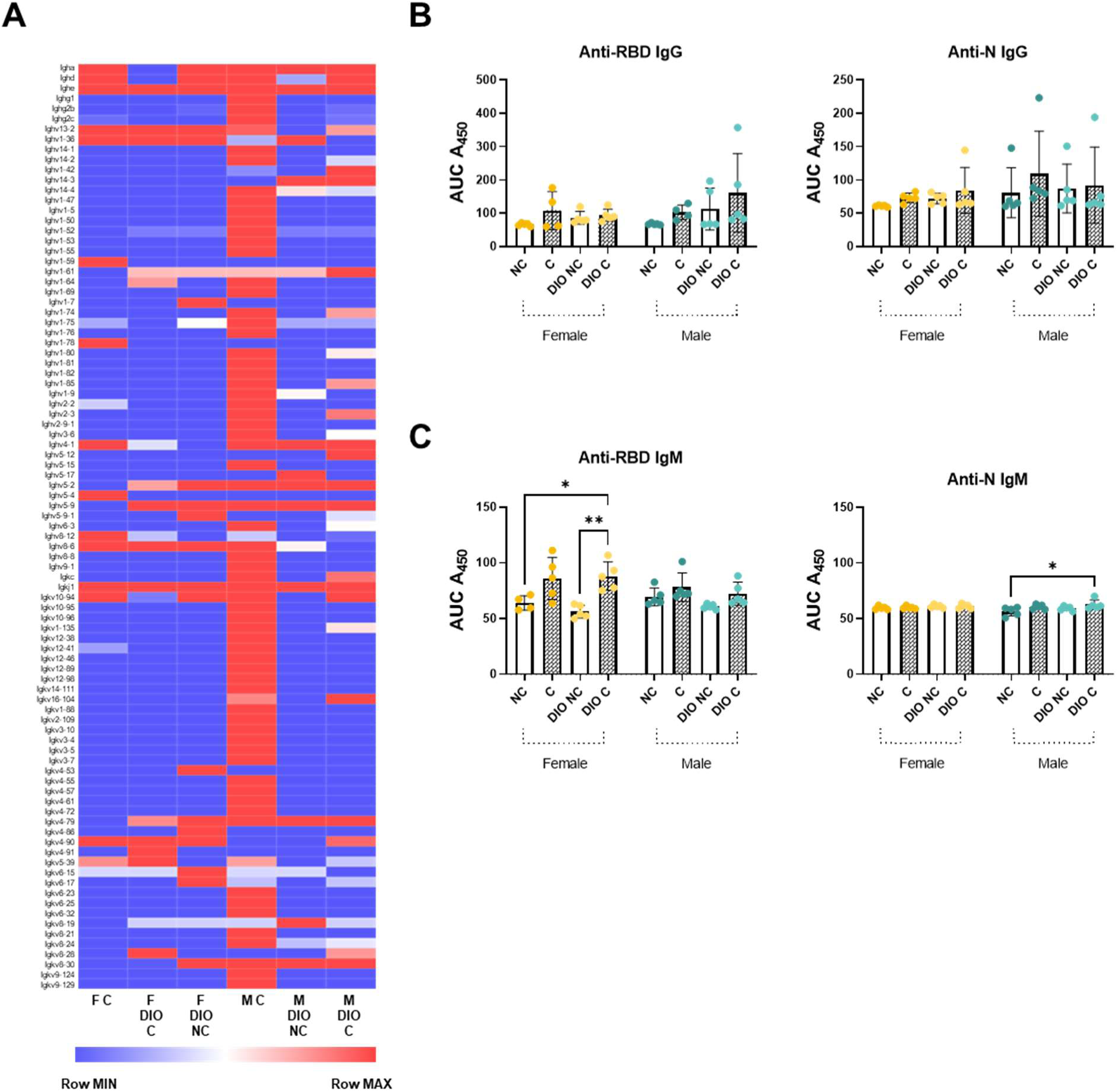
Antibody gene expression due to SARS-CoV-2 in the lungs of DIO K18-hACE2 mice. Antibody gene counts from lung RNA of mice challenged with SARS-CoV-2 (A). Anti-RBD and anti-nucleocapsid (N) IgG and IgM measured in serum of K18-hACE2 mice (B). (*P* value: * < 0.0200; ** = 0.0454)

## Discussion

Despite vaccines, antibodies, and small molecule therapeutics, SARS-CoV-2 continues to infect individuals driving continuation of the COVID-19 pandemic. Due to the heterogenous nature of COVID-19 cases, it is important to consider the host factors and responses that predispose to severe infection or death. Obesity and Type 2 Diabetes (T2DM) have long been appreciated as co-morbidities for infectious diseases, yet very little controlled experimental data has illuminated how these comorbidities interplay with COVID-19. Additionally, sex-specific responses to COVID-19 are obvious in the diversity of disease manifestation (86). To begin the effort to explore these conditions in preclinical models, we aimed to develop a comorbidity model for COVID-19 based on the previously implicated model of diet induced obesity and T2DM (Fig. 1). Utilizing our model, we designed experiments to evaluate sex and DIO as central variables. We utilized the Alpha variant of SARS-CoV-2 which we previously identified as having high virulence compared to ancestral strains (55). K18-hACE2 mice with DIO showed that disease as well as morbidity and mortality were affected very little by the addition of DIO in males; however, the DIO condition greatly affected females (Fig. 2) and resulted in 100-fold higher viral RNA burden in the lung (Fig. 3). In order to characterize the augmented host response to viral challenge we utilized RNAseq analysis of the lung tissue to analyze the pathogen-specific airway responses to the presence of virus (Fig. 4). Pathway analysis using GO term and Ingenuity Pathway analyses illuminated more sex- and diet-specific responses. Notably, it appeared that DIO non-challenged male mice develop preexisting T cell gene expression signatures in the lungs suggestive of T cell infiltration that is decreased after viral challenge (Fig. 6A). DIO seemed to hinder B cell responses in the lungs of challenged male mice (Fig. 6-8) indicating that while DIO did not affect the development of morbidity in males, it did alter the host response to infection. In female mice, distinctive gene expression profiles were observed between normal diet and DIO mice. Specifically, challenged female mice have low gene expression profiles corresponding to IPA’s Coronavirus Pathogenesis Pathway, while the addition of the DIO condition enhances many of the pathway’s genes (Fig. 6). Our data suggest that female K18-hACE2-mice on normal diets have lower viral RNA burden and decreased inflammatory responses, and DIO impairs the clearance of virus in the lung which results in their enhanced morbidity (Fig. 2-3). Collectively these data begin to shed light on the effects of DIO on COVID-19.

To the best of our knowledge this is the first study in K18-hACE2-mice to evaluate DIO and sex utilizing transcriptomic analysis to better understand SARS-CoV-2 VOC-specific host responses. A previous MedRxiv study utilized a mouse-adapted strain of SARS-CoV-2 to challenge DIO C57Bl6 mice to measure the protective efficacy of human convalescent serum treatment (87). One correspondence describes a small study where DIO mice were challenged with SARS-CoV-2 and the authors reported increased lung pathology and interferon responses (27). However, the study did not utilize sex comparisons nor analyzed transcriptomic responses. The high-fat high-carbohydrate diet has been used to evaluate the combined effects of the “Western Diet” and COVID-19 disease in Syrian hamsters (88). Comparable to what we observed in mice, Western

Diet-affected hamsters had increased weight loss, lung pathology, and delayed viral clearance after challenge. Our study does have some caveats that warrant discussion. In our experiment, we only evaluated one SARS-CoV-2 challenge strain, Alpha, and there have now been three VOC strain surges (Beta, Delta, Omicron) since Alpha was dominantly circulating. In additional studies since then, we have observed enhanced airway inflammation due to challenge with the Delta variant (68). We anticipate that different strains would result in variable host responses to what was identified using Alpha; however, additional studies will need to be performed. Another caveat is that only one challenge dose was evaluated (1,000 PFU). If lower or higher challenge doses were to be studied, we would expect to have either shorter or longer time to morbidity during which host response profiles may further develop or remain hidden due to the disease timeline. Finally, our study focused on defining transcriptomic responses to characterize the altered host responses to SARS-CoV-2 challenge. We did not analyze specific cell populations through cell isolation and flow cytometry, nor did we evaluate potential mechanisms responsible for this co-morbidity.

SARS-CoV-2 infection in humans is generally heterogenous in symptomology, however, increased susceptibility to severe infection requiring hospitalization are common across patients with T2DM metabolic disease and increased adiposity (obesity) (89). The role of elevated glucose and fat accumulation downstream of metabolic dysfunction has been shown in other settings to disturb cellular signaling cascades, promote cytokine synthesis and secretion (leading to hyperinflammation), and increase oxidative stress through enhancement of reactive oxygen species (ROS)(90). However, the mechanism by which elevated glucose and adiposity enhance the severity of SAR-CoV-2 infection is still unclear. Our data, along with human RNA-Seq analysis from T2DM samples, demonstrates the altered immune system activation in response to infection largely through changes in antibody response. Indeed, human enrichment and GO term analysis in our mouse model compared with lung epithelium from T2DM human patients shows TNF and IL-17 signaling to be highly enriched along with several other genes involved in the antibody response (91). As demonstrated in our findings, the inflammatory signature of T2D hosts is further activated following infection supporting the hypothesis that the preexisting inflammatory state of patients with T2DM and/or obesity contributes to the severity of infection driving predisposition to negative outcomes (92). We also saw that T cell populations were augmented across our study groups with male DIO challenged mice demonstrating the most noticeable decrease in T cell subtypes. In severe human COVID-19 cases, it has been demonstrated that lymphocytopenia (reduced T cell counts) is probably associated with CD4+ and CD8+ T cell exhaustion (93–99). This may be another contributor towards SARS-CoV-2’s severity in T2DM. Lastly, our RNA-Seq data in normal diet challenge mice compared with normal diet infected patient samples also demonstrates overlap in gene expression profiles, highlighting the relevance and use of the k18-hACE2 model (100, 101). We believe that our model demonstrates some of the pathology seen in human cohorts (summarized in Fig. 9). This overlap in human and mouse data is encouraging as modeling SAR-CoV-2 in the laboratory setting is critical for understanding the molecular mechanisms at work.

**Figure 9:**
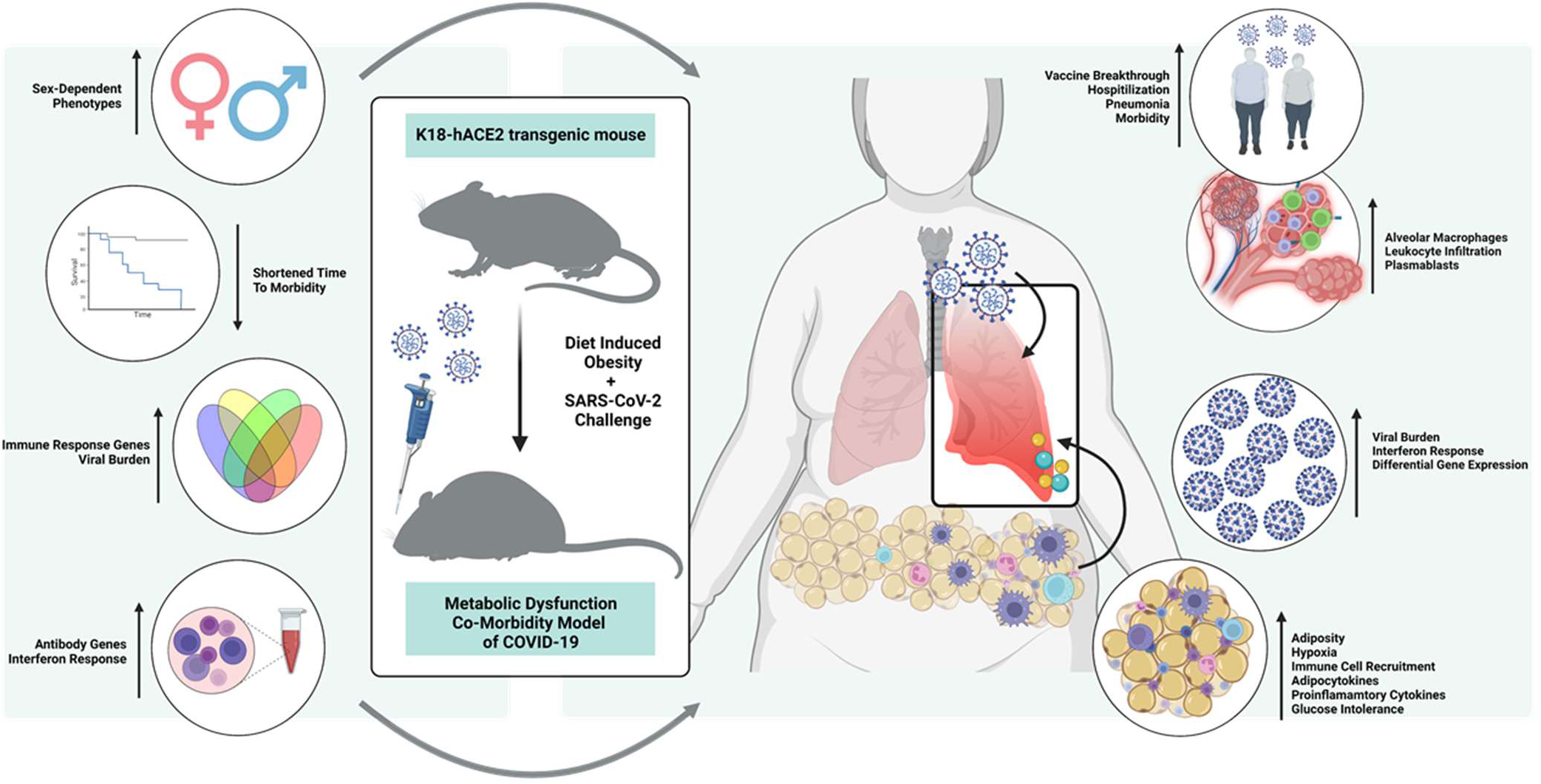
The DIO-COVID-19 mouse model hypothesis: translating mouse data to clinical phenotypes. Our data suggests DIO promotes sex-dependent phenotypes including unique differential gene expression and a shortened time to morbidity when challenged with SARS-CoV-2. Gene expression analysis parallels the changes in viral RNA burden through elevation of immune response genes and the interferon response. As persons with metabolic disease are more likely to succumb to severe SARS-CoV-2 infection and experience vaccine breakthrough, understanding the altered adaptive response is necessary for identifying therapeutic targets. We hypothesize thar hyperglycemia and SARS-CoV-2 infection have a synergistic effect in altering the lung transcriptome in comparison to viral infection alone. Perhaps it is this synergistic effect that is the central driver of the maladapted immune response.

The T2DM-COVID-19 mouse model has allowed us to observe responses to SARS-CoV-2 challenge augmented by both obesity and sex. It is apparent that DIO affects female mice and enhances viral virulence; however, it is not clear how to best ameliorate this issue using therapeutic interventions. In a prior study (data unpublished), we evaluated treatment with Baricitinib, a JAK inhibitor that could dampen inflammation caused by SARS-CoV-2. We hypothesized that decreasing inflammation would improve survival outcomes of SARS-CoV-2-challenged mice; however, the drug did not improve survival of SARS-CoV-2 challenged mice (data not shown). This data leads us to believe that there are still molecular events that underpin overall inflammation as well as a dynamic viral clearance timeline that need to be adjusted to improve protection. Comparing that study with our DIO model, it is clear we need to profile the cell populations that respond to each phase of viral infiltration and identify if they are ineffective at stopping disease progression. Furthermore, we propose that this DIO model can be used to evaluate differences in vaccine-induced immunity among comorbid groups which is rarely evaluated. We plan to continue our utilization of the novel DIO-COVID-19 mouse model to uncover therapeutic strategies to improve disease and survival outcomes in diverse persons infected with SARS-CoV-2.

## Supporting information

Supplemental Expression Browser Data for RNAseq

## Acknowledgements

This project was executed with support from the Vaccine Development Center at the West Virginia University Health Sciences Center. F.H.D. and the VDC are supported by the Research Challenge Grant no. HEPC.dsr.18.6 from the Division of Science and Research, WV Higher Education Policy Commission. We thank BioRender for the use of their software to create the final figure. Lastly, we thank Drs. Laura Gibson and Clay Marsh for supporting this and the lab’s additional COVID-19 research efforts.

## Author contributions

Studies were designed by FHD, HAC, KSL, BPR, JRB, AMH. All authors contributed to the execution of the studies. MTW and IM prepared and provided titred viral stocks of SARS-CoV-2 for challenge. Animal health checks, necropsy, and tissue processing were performed by FHD, TYW, BPR, KSL, JRB, OAM, AMH, and HAC. Viral RNA qPCR was performed by HAC and OAM. Serological analysis was executed by KSL, BPR, and NAR. Luminex cytokine assays were completed by BPR. MSD assays were performed by KSL. Data was analyzed by KSL, BPR, HAC, and FHD. All authors contributed to the writing and revision of this manuscript.

